# Cadherin-7 mediates proper neural crest cell-placodal neuron interactions during trigeminal ganglia assembly

**DOI:** 10.1101/434613

**Authors:** Chyong-Yi Wu, Lisa A. Taneyhill

## Abstract

The cranial trigeminal ganglia play a vital role in the peripheral nervous system through their relay of sensory information from the vertebrate head to the brain. These ganglia are generated from the intermixing and coalescence of two distinct cell populations: cranial neural crest cells and placodal neurons. Trigeminal ganglia assembly requires the formation of cadherin-based adherens junctions within the neural crest cell and placodal neuron populations; however, the molecular composition of these adherens junctions is still unknown. Herein, we aimed to define the spatio-temporal expression pattern and function of Cadherin-7 during early chick trigeminal ganglia formation. Our data reveal that Cadherin-7 is expressed exclusively in migratory cranial neural crest cells and is absent from trigeminal neurons. Using molecular perturbation experiments, we demonstrate that modulation of Cadherin-7 in neural crest cells influences trigeminal ganglia assembly, including the organization of neural crest cells and placodal neurons within the ganglionic anlage. Moreover, alterations in Cadherin-7 levels lead to changes in the morphology of trigeminal neurons. Taken together, these findings provide additional insight into the role of cadherin-based adhesion in trigeminal ganglia formation, and, more broadly, the molecular mechanisms that orchestrate the cellular interactions essential for cranial gangliogenesis.

## INTRODUCTION

The cranial ganglia of the peripheral nervous system perform crucial sensory functions, including somatosensation and innervation of specific organs such as the heart and lungs. The trigeminal ganglion (cranial nerve V) is responsible for the former, mediating sensations of pain, touch, and temperature in the face and innervating the sensory apparti of the muscles of the eye and upper and lower jaws. The collective intermixing and condensation of two embryonic cell populations, neural crest cells and neurogenic placode cells, is required to assemble the cranial ganglia (Breau and Schneider-Maunoury, 2015; Hamburger, 1961; Saint-Jeannet and Moody, 2014; Steventon et al., 2014). Neural crest cells arise from the dorsal region of the developing neural folds, undergo an epithelial-to-mesenchymal transition, and migrate to stereotypical destinations depending upon their axial level of origin and the molecular cues received from the extracellular environment (Bronner and Simoes-Costa, 2016; Duband et al., 2015; Gouignard et al., 2018; Simoes-Costa and Bronner, 2015; Taneyhill and Schiffmacher, 2017). Neurogenic placode cells originate as paired epidermal thickenings at distinct rostral-caudal positions in the vertebrate head. These cells delaminate from the surface ectoderm (and while doing so begin differentiating) and then migrate through the cranial mesenchyme, where they will eventually coalesce with neural crest cells to form the cranial ganglia (Baker and Bronner-Fraser, 2001; Jidigam and Gunhaga, 2013; Smith et al., 2015; Steventon et al., 2014). Prior studies have highlighted the reciprocal nature of neural crest cell-placodal neuron interactions, providing a working hypothesis in which neural crest cells act as a scaffold to integrate all of the placodal neurons such that one ganglion forms, while placodal neurons, in turn, facilitate neural crest cell condensation (D’Amico-Martel and Noden, 1983; Hamburger, 1961; Shiau et al., 2008). The interactions between neural crest cells and placodal neurons are also highly dynamic, as these cells exhibit a “chase and run” behavior (Theveneau et al., 2013), with neural crest cells forming favorable pockets, or corridors, upon which placodal neurons prefer to migrate versus the less permissive mesoderm (Freter et al., 2013). As such, both transient and more stable intercellular interactions must occur between these two distinct cell types during their migration and coalescence to form a tightly adhered tissue.

Previous work has identified components of cadherin-based cellular adherens junctions, the levels of which must be tightly controlled to allow for proper intercellular interactions and gangliogenesis. In the chick, placodal neurons express N-cadherin, and signaling between Slit1, secreted by neural crest cells, and Robo2, the Slit1 receptor on the surface of placodal neurons, regulates N-cadherin levels, likely through a post-translational mechanism (Shiau et al., 2008; Shiau and Bronner-Fraser, 2009). Moreover, N-cadherin knockdown phenocopies depletion of Robo2, leading to more dispersed placodal neurons and, ultimately, defects in ganglion condensation (Shiau and Bronner-Fraser, 2009). The presence of *Cadherin-7* transcripts in the chick embryo was reported over 20 years ago using whole-mount *in situ* hybridization, which revealed *Cadherin-7* expression in early migrating cranial neural crest cells (Hamburger-Hamilton stage 10 (HH10)) as well as in the forming trigeminal ganglion (HH18) (Nakagawa and Takeichi, 1995). Long-term overexpression of Cadherin-7 in chick trunk neural crest cells was shown to only abrogate migration of neural crest cells along the dorsolateral pathway (taken by melanocyte precursors) but did not inhibit neural crest cell differentiation (Nakagawa and Takeichi, 1998). The specific effect on melanocytes was ascribed to the timing at which the viral construct expressing Cadherin-7 achieved maximal levels (Nakagawa and Takeichi, 1998), and thus a functional role for Cadherin-7 in the neural crest was still not fully appreciated. More recent work revealed the presence of αN-catenin in migratory cranial neural crest cells, with perturbations in αN-catenin impacting trigeminal ganglia assembly, in part, through changes in the placodal neuron contribution to the ganglia and the level of Cadherin-7 in neural crest cells (Wu et al., 2014). No studies to date, however, have documented the spatio-temporal expression pattern of Cadherin-7 protein in chick migratory cranial neural crest cells during early gangliogenesis, nor investigated the role of Cadherin-7 in neural crest cells that form the trigeminal ganglia.

To this end, we have undertaken studies to define the distribution and function of Cadherin-7 in migratory neural crest cells during early chick trigeminal gangliogenesis. Our data show that Cadherin-7 is expressed solely in migratory cranial neural crest cells as the trigeminal ganglia form. To address Cadherin-7 function, we depleted and overexpressed Cadherin-7 in migratory neural crest cells and evaluated embryos for effects on trigeminal ganglia assembly. In each instance, we noted alterations in the distribution of neural crest cells and trigeminal neurons, along with abnormal trigeminal neuron morphology. Collectively, these results suggest that Cadherin-7 plays an important role in the migratory cranial neural crest cell population to permit correct trigeminal ganglia formation in the early chick embryo.

## RESULTS

### Cadherin-7 is expressed in migratory cranial neural crest cells contributing to the

We documented the spatio-temporal expression pattern of Cadherin-7 protein throughout early stages of chick trigeminal gangliogenesis (HH12-HH17) using confocal microscopy. In keeping with a previously published report on *Cadherin-7* transcripts in the chick head (Nakagawa and Takeichi, 1995) and our prior study (Wu et al., 2014), we noted Cadherin-7 protein in migratory cranial neural crest cells (Fig. 1A-D) identified by labeling with an antibody to HNK-1 (Bronner-Fraser, 1986) (Fig. 1A’-D’, arrows) throughout all stages examined (HH13-16 shown, identical results observed for HH12 and HH17). Cadherin-7 is primarily observed on the plasma membrane of neural crest cells, co-localizing with the cell surface HNK-1 neural crest cell marker (Fig. 1A’-D’, arrows). Moreover, Cadherin-7 protein is observed in the neural tube (Fig. 1A-D, *). In contrast, Cadherin-7 is not detected in the trigeminal placode precursors residing in the surface ectoderm nor in trigeminal neurons, here labeled with Annexin A6 (Fig. 1A’-D’, arrowheads), as only placodal neurons express Annexin A6 at these stages of development (Shah and Taneyhill, 2015). The cranial mesenchyme is devoid of Cadherin-7, as noted previously (Nakagawa and Takeichi, 1995). Thus, Cadherin-7 is expressed exclusively in migratory cranial neural crest cells throughout early trigeminal gangliogenesis

**Figure 1.**
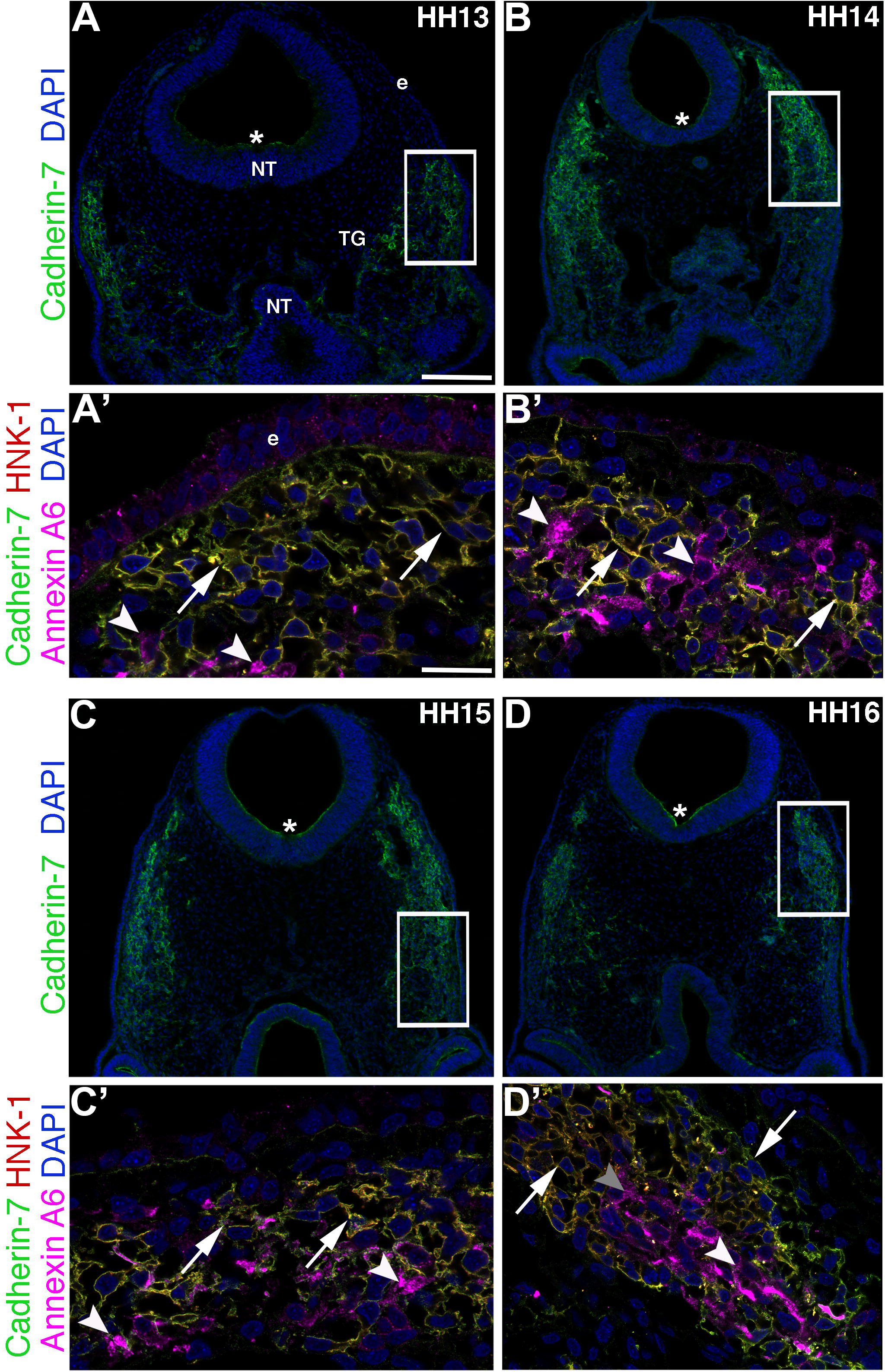
Cadherin-7 protein is observed in migratory neural crest cells contributing to the cranial trigeminal ganglia. Representative transverse sections taken at the axial level of the forming trigeminal ganglia over four chick embryo stages (HH13-16) followed by immunohistochemistry for Cadherin-7 (green), HNK-1 (red, labels neural crest cells), and Annexin A6 (purple, labels placodal neurons). (A-D) Lower magnification images show the entire transverse section and reveal Cadherin-7 immunoreactivity. Higher magnification images (A’-D’) of the boxed area in (A-D) show Cadherin-7- and HNK-1-double-positive neural crest cells at all stages (arrows), whereas Annexin A6-positive placodal neurons are devoid of Cadherin-7. DAPI (blue) labels cell nuclei. Ectoderm, neural tube, and trigeminal ganglion are denoted by e, NT, and TG, respectively. Scale bar in (A) is 100μm and applicable to (B-D), while scale bar in (A’) is 5μm and applicable to (B’-D’).

### Reduced levels of Cadherin-7 alter trigeminal ganglia assembly through effects on neural crest cells and placodal neurons

To assess the functional role of Cadherin-7 in neural crest cells contributing to the trigeminal ganglia, we performed morpholino (MO)-mediated knockdown of Cadherin-7 using a translation-blocking MO that targets the *Cadherin-7* 5’ UTR and initial coding region (see Materials and Methods). As a control, we designed a five bp mismatch Cadherin-7 MO that does not block *Cadherin-7* translation. MO efficacy was confirmed by immunoblotting for Cadherin-7 protein in lysates prepared from electroporated, dissected trigeminal ganglia, as in (Shah et al., 2017), which resulted in an approximate 50% reduction in Cadherin-7 protein levels in tissue possessing the Cadherin-7 MO versus the control MO (Supp. Fig. 1, n = 2;).

We next determined whether knockdown of Cadherin-7 affects neural crest cells and placodal neurons contributing to the trigeminal ganglia. Premigratory cranial neural crest cells were unilaterally electroporated with the Cadherin-7 or control MO, and embryos were allowed to grow to HH15-HH16 followed by immunohistochemistry on cranial transverse sections using molecular markers to label neural crest cells (HNK-1) and placodal neurons (Tubb3), the latter of which only labels placode cell-derived neurons at this stage of development (Moody et al., 1989; Shiau et al., 2008), as neural crest cells differentiate much later (D’Amico-Martel and Noden, 1980; Steventon et al., 2014). Introduction of the control MO (Fig. 2A) into migratory neural crest cells did not affect the distribution of neural crest cells and trigeminal neurons, as assessed by HNK-1 (Fig. 2B) and Tubb3 (Fig. 2C) immunohistochemistry (n = 6 and n = 5 embryos, respectively), nor their ability to coalesce together (Fig. 2D, D’, arrows and arrowheads). Depletion of Cadherin-7 through introduction of the Cadherin-7 MO (Fig. 2E), however, impacted both the neural crest cell and trigeminal neuron populations in the ganglionic anlage. Neural crest cells did not organize correctly to form the typical morphology of the ganglion (Fig. 2F, H’, arrows; n = 13/13 embryos; compare to Fig. 2B, D’, arrows). Interestingly, trigeminal neurons were also affected, with these cells possessing fewer neuronal projections and thus appearing round instead of bipolar (Fig. 2G, H’, arrowheads; n = 13/13 embryos) compared to the morphology adopted by trigeminal neurons in control MO-treated embryos (Fig. 2C, D’, arrowheads). In addition, trigeminal neurons tended to aggregate together in small clusters or groups (Fig. 2G, H’, arrowheads).

**Figure 2.**
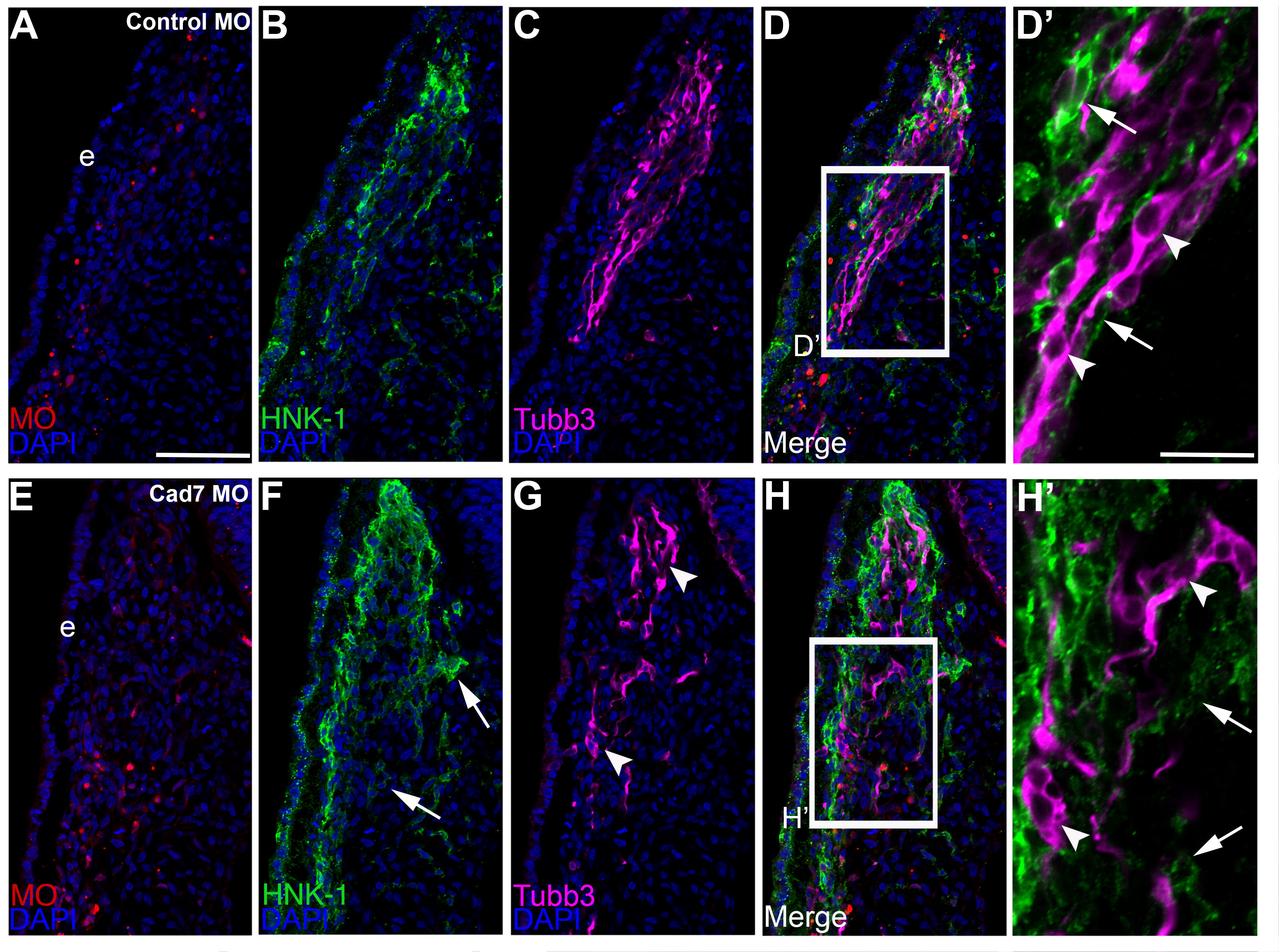
Morpholino-mediated depletion of Cadherin-7 from migratory cranial neural crest cells alters the distribution of neural crest cells and placodal neurons within the forming trigeminal ganglia. Representative transverse sections taken at the axial level of the forming trigeminal ganglia after electroporation of a 5 bp mismatch control Cadherin-7 morpholino (Control MO, A-D′) or Cadherin-7 MO (Cad7 MO, E-H′) into premigatory neural crest cells at the 3ss followed by immunohistochemistry for HNK-1 (green) and Tubb3 (purple). (D’, H’) Higher magnification images of the boxed regions in (D, H). (A-D’) A typical trigeminal ganglion containing the control MO in the neural crest (A) exhibits a stereotypical “tear drop” morphology in section at this axial level due to the coalescence of neural crest cells (B, D) with placodal neurons (C, D). At higher magnification (D’), HNK-1-positive neural crest cells (arrows) surround Tubb3-positive placodal neurons (arrowheads), many of which are already becoming bundled and adopting the bipolar morphology associated with neuronal maturation. Conversely, a trigeminal ganglion containing the Cadherin-7 MO in the neural crest (A) possess an abnormal morphology due to the position of the neural crest cells (F, H, arrows) and placodal neurons (G, H, arrowheads) within the anlage. At higher magnification (H’), it is apparent that neural crest cells still surround the placodal neurons (arrows), but the shape adopted by the placodal neurons is aberrant, with neurons appearing round (arrowheads). DAPI (blue) labels cell nuclei. e, ectoderm. Scale bar in (A) is 67μm and applicable to (B-H), while scale bar in (D’) is 20μm and applicable to (H′).

To rule out potential indirect effects on neural crest cells that could be causing these phenotypes during ganglia assembly, such as changes in cell proliferation or cell death, we next performed phospho-histone H3 (PHH3) immunohistochemistry as well as a TUNEL assay, respectively. Upon counting PHH3- or TUNEL-positive cells and comparing the contralateral control and MO-treated sides, we noted no statistically significant difference in either cell proliferation (Supp. Fig. 2A-D, arrows; 24 +/− 1 PHH3-positive cells for the control MO-treated side, 23 +/− 1 PHH3-positive cells for the contralateral side, p = 0.84; 24 +/− 1 PHH3-positive cells for the Cadherin-7 MO-treated side, 25 +/− 1 PHH3-positive cells for the contralateral side, p = 0.71) or cell death (Supp. Fig. 2E-H, arrows; 31 +/− 2 TUNEL-positive cells for the control MO-treated side, 30 +/− 3 TUNEL-positive cells for the contralateral side, p = 0.63; 24 +/− 2 TUNEL-positive cells for the Cadherin-7 MO-treated side, 23 +/− 2 TUNEL-positive cells for the contralateral side, p = 0.71). As such, the overall organization of the trigeminal ganglion appeared abnormal upon Cadherin-7 knockdown in neural crest cells due to effects on both migratory neural crest cells and placodal neurons unrelated to changes in cell proliferation or cell death.

Given the observed phenotypes in tissue sections, we next examined the distribution of neural crest cells and placodal neurons within the context of the entire trigeminal ganglion by performing whole-mount immunohistochemistry for HNK-1 and Tubb3, respectively. Lateral views of whole embryo heads at HH15-HH16 obtained by confocal microscopy revealed the normal distribution of neural crest cells and placodal neurons within the trigeminal ganglion in the presence of the control MO (Fig. 3A-D, n = 10/10 embryos). In these images, the bundling of placodal neurons, and condensation with migratory neural crest cells that closely localized with placodal neurons, generated the stereotypical structure of the trigeminal ganglion that is revealed through the Tubb3-positive immunoreactivity of the placodal neurons (Fig. 3C, D, arrowheads). The morphology of the trigeminal ganglion is perturbed, however, upon MO-mediated knockdown of Cadherin-7 (Fig. 3E-H, n = 11/13 embryos). From these experiments, it is apparent that Cadherin-7-depleted neural crest cells still localized to the anlage, allowing them to intermingle with trigeminal neurons (compare Fig. 3F to Fig. 3B). Even with this seemingly correct localization, though, trigeminal neurons no longer bundled together correctly, leading to the appearance of a disorganized, less condensed trigeminal ganglion (compare Fig. 3G, H, arrowheads, to Fig. 3C, D, arrowheads). Collectively, these data provide evidence for a role for Cadherin-7 in neural crest cells during trigeminal ganglion assembly.

**Figure 3.**
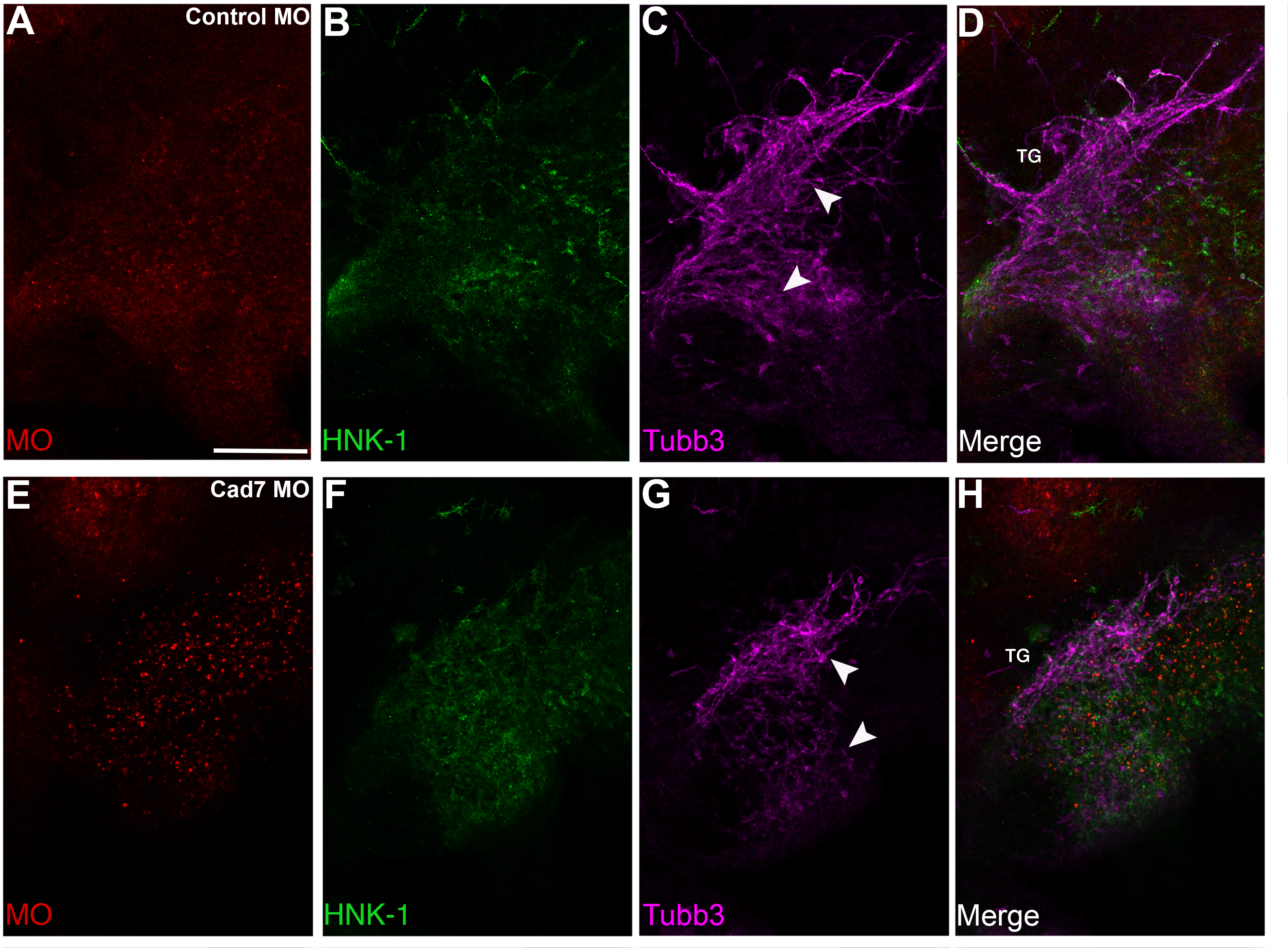
Cadherin-7 depletion in migratory neural crest cells alters the gross morphology of the trigeminal ganglion. Representative lateral views (optical section) of the forming trigeminal ganglion in an HH15 chick head after electroporation of a 5 bp mismatch control Cadherin-7 MO (Control MO, A-D) or Cadherin-7 MO (Cad7 MO, E-H) at the 3ss, followed by whole-mount immunohistochemistry for HNK-1 (green) and Tubb3 (purple). Merge images are shown in (D, H). Trigeminal ganglia electroporated with the control MO in neural crest cells (A) exhibit a condensed, organized morphology, with neural crest cells (B) associating with placodal neurons that are forming nerve bundles (C, arrowheads). Those trigeminal ganglia electroporated with the Cadherin-7 MO in the neural crest (E) possess neural crest cells that migrate to the anlage (F) but exhibit less bundling of placodal neurons (G, arrowheads), leading to an aberrant ganglion shape relative to control. TG, trigeminal ganglion. Scale bar in (A) is 200μm and applies to all images.

### Overexpression of Cadherin-7 negatively affects trigeminal ganglia assembly by impacting both neural crest cells and placodal neurons

To further elucidate the function of Cadherin-7 in migratory cranial neural crest cells, we overexpressed Cadherin-7 and evaluated effects on the neural crest and placodal neuron populations forming the trigeminal ganglia. Cadherin-7 protein levels were increased by 200% over control, as assessed by immunoblotting for Cadherin-7 in lysates prepared from electroporated, dissected trigeminal ganglia possessing the control vector (pCIG) versus the Cadherin-7 expression construct (pCIG-Cad7, contains an IRES-GFP cassette to label electroporated cells; see Materials and Methods for details) (Supp. Fig. 3, n = 2). Next, we evaluated how augmented Cadherin-7 protein levels in neural crest cells affect trigeminal ganglia formation. To this end, we performed similar neural crest cell electroporation experiments and analyzed tissue sections (Fig. 4) and whole embryo heads (Fig. 5) for changes in neural crest cells and/or placodal neurons contributing to the trigeminal ganglia. In the presence of the control pCIG vector (Fig. 4A), we noted no alterations in migratory neural crest cells (Fig. 4B, n = 8/8 embryos) or placodal neurons (Fig. 4C, n = 8/8 embryos). These results indicate that the trigeminal ganglia assembled normally (Fig. 4D) and that the electroporation technique did not affect its formation. Cadherin-7 overexpression in neural crest cells, however, negatively impacted trigeminal ganglia assembly. Neural crest cells expressing increased levels of Cadherin-7 (Fig. 4E) did not associate with one another (and with placodal neurons) properly to generate the morphology of the ganglion that is typically observed upon sectioning (Fig. 4F, H’, arrows; n = 13/15 embryos; compare to Fig. 4B, D’, arrows). In turn, placodal neurons were also affected, exhibiting, in some instances, a more round morphology, as opposed to the bipolar shape that these neurons normally possess, and were generally misshapen (Fig. 4G, H’, arrowheads; n = 21/22 embryos; compare to Fig. 4C, D’, arrowheads).

**Figure 4.**
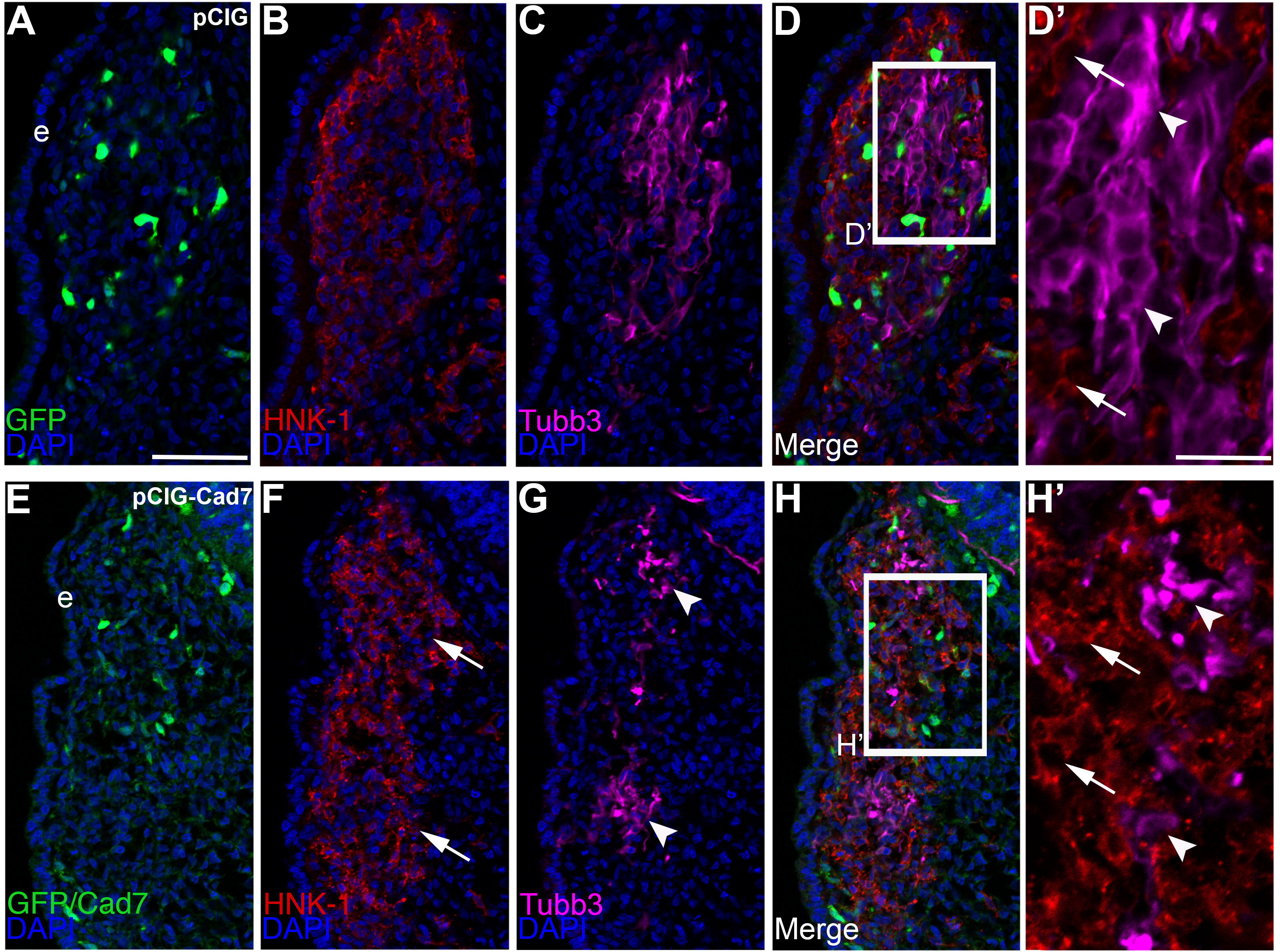
Overexpression of Cadherin-7 in migratory cranial neural crest cells alters the distribution of neural crest cells and placodal neurons within the forming trigeminal ganglia. Representative transverse sections taken at the axial level of the forming trigeminal ganglia after electroporation of the pCIG control vector (pCIG, A-D′) or the pCIG-Cadherin-7 vector (pCIG-Cad7, E-H′) into premigatory neural crest cells at the 3ss followed by immunohistochemistry for HNK-1 (red) and Tubb3 (purple). The pCIG vector contains an IRES-GFP cassette to label electroporated cells. (D’, H’) Higher magnification images of the boxed regions in (D, H). (A-D’) A trigeminal ganglion containing the control pCIG vector in the neural crest (A) possesses a stereotypical “tear drop” morphology that is noted in section at this axial level due to the coalescence of neural crest cells (B, D) with placodal neurons (C, D). At higher magnification (D’), HNK-1-positive neural crest cells (arrows) form corridors around Tubb3-positive placodal neurons (arrowheads), many of which are elaborating neuronal protrusions indicative of neuronal maturation. On the other hand, neural crest cells with elevated levels of Cadherin-7 protein (E) do not distribute correctly in the ganglionic anlage (F, H, arrows), and placodal neurons also localize incorrectly (G, H, arrowheads). At higher magnification (H’), neural crest cells are noted around the placodal neurons (arrows), but placodal neuron morphology is abnormal, with neurons appearing round and/or misshapen (arrowheads). DAPI (blue) labels cell nuclei. e, ectoderm. Scale bar in (A) is 50μm and applicable to (B-H), while scale bar in (D’) is 20μm and applicable to (H′).

**Figure 5.**
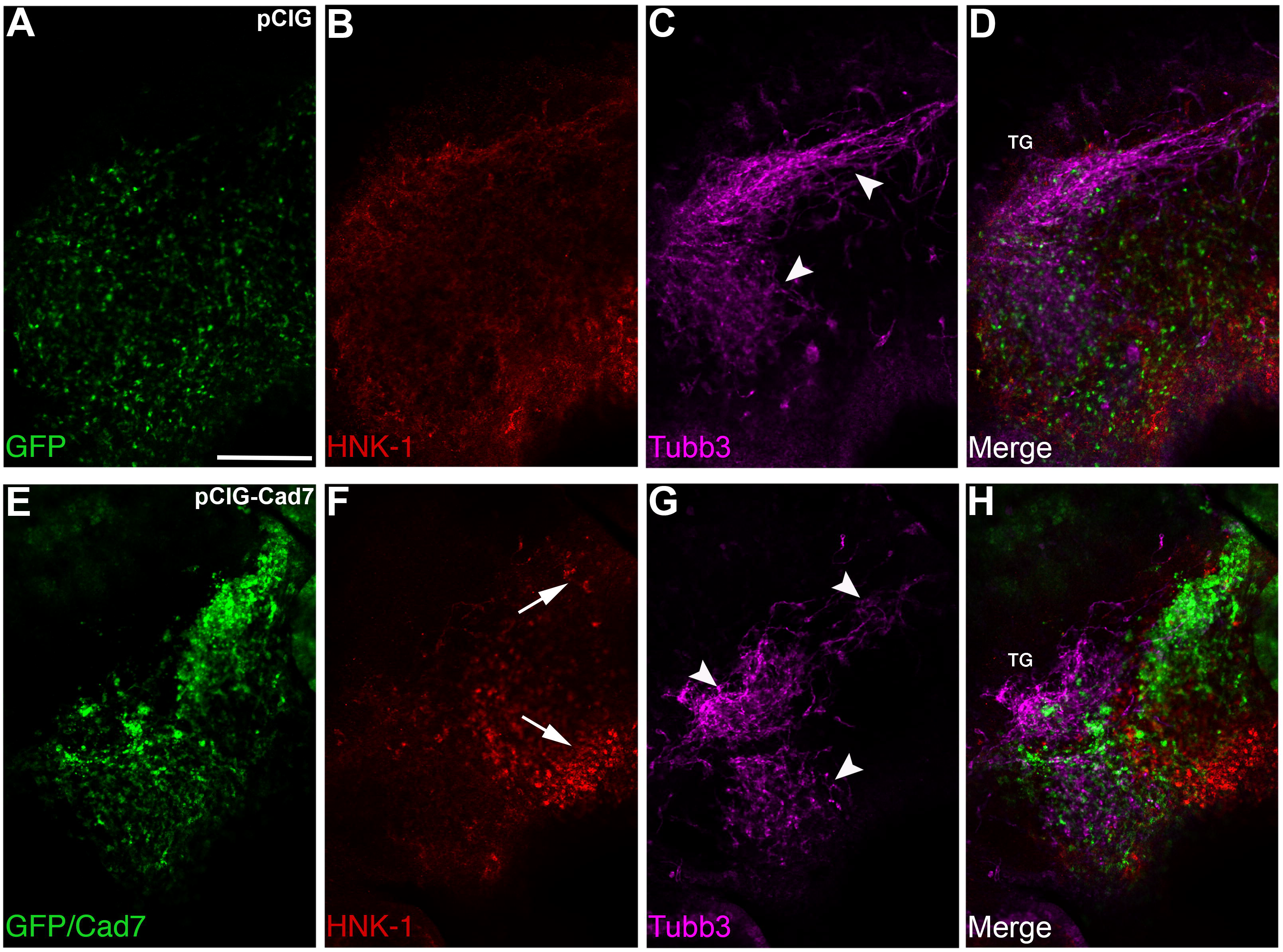
Elevated levels of Cadherin-7 in migratory neural crest cells alter the gross morphology of the trigeminal ganglia. Representative lateral views (optical section) of the forming trigeminal ganglion in an HH15 chick head after electroporation of the pCIG control vector (pCIG, A-D) or the pCIG-Cadherin-7 vector (pCIG-Cad7, E-H) at the 3ss, followed by whole-mount immunohistochemistry for HNK-1 (red) and Tubb3 (purple). Merge images are shown in (D, H). Trigeminal ganglia electroporated with the control pCIG vector in neural crest cells (A) exhibit a condensed, organized morphology, with neural crest cells (B) associating with placodal neurons that are forming nerve bundles (C, arrowheads). Those trigeminal ganglia electroporated with pCIG-Cad7 in the neural crest (E) possess neural crest cells that migrate to the anlage but appear to aggregate together, thus altering their general distribution in the anlage (F, arrows). Placodal neurons are also affected, exhibiting less bundling and an overall disorganized phenotype (G, arrowheads). Together, this leads to an abnormal ganglion shape relative to control. TG, trigeminal ganglion. Scale bar in (A) is 200μm and applies to all images.

To confirm that the observed phenotypes were not due to non-specific effects on cell proliferation or death, we conducted phospho-histone H3 immunohistochemistry and a TUNEL assay, respectively. We noted no statistically significant change in either cell proliferation (Supp. Fig. 4A-D, arrows; 31 +/− 1 PHH3-positive cells for the pCIG control-treated side, 30 +/− 1 PHH3-positive cells for the contralateral side, p = 0.82; 26 +/− 1 PHH3-positive cells for the pCIG-Cad7-treated side, 26 +/− 1 PHH3-positive cells for the contralateral side, p = 0.72) or cell death (Supp. Fig. 4E-H, arrows; 22 +/− 2 TUNEL-positive cells for the pCIG control-treated side, 21 +/− 2 TUNEL-positive cells for the contralateral side, p = 0.71; 31 +/− 2 TUNEL-positive cells for the pCIG-Cad7-treated side, 30 +/− 3 TUNEL-positive cells for the contralateral side, p = 0.78) in the presence of the control vector or Cadherin-7 overexpression construct. Collectively, these results further confirm that Cadherin-7 expression in neural crest cells is important for appropriate trigeminal ganglia assembly and that the noted phenotypes are not caused by changes in cell proliferation or cell death.

Given these results in tissue sections, we then analyzed whole embryo heads from HH15-HH16 embryos by confocal microscopy following immunostaining for HNK-1 and Tubb3, as we had done previously with our MO-electroporated embryos (Fig. 5). Introduction of the pCIG control vector (Fig. 5A) into migratory neural crest cells revealed no effects on neural crest cell-placodal neuron interactions and, ultimately, trigeminal gangliogenesis (Fig. 5A-D, n = 7/7 embryos), as exemplified by Tubb3 immunoreactivity and the formation of a ganglion possessing a bi-lobed structure (Fig. 5C, arrowheads). Cadherin-7 overexpression, however, disrupted trigeminal ganglia assembly, as evidenced by the presence of a disorganized ganglion in which placodal neurons did not properly condense with neural crest cells (Fig. 5E-H, n = 7/7 embryos). While HNK-1-positive neural crest cells migrated and localized to the ganglionic anlage, their distribution was noticeably different than that observed with the pCIG control vector, with neural crest cells aggregating together (compare Fig. 5E, F, arrows, to Fig. 5A, B). Furthermore, trigeminal neurons did not form stereotypical bundles and appeared more disorganized within the anlage (compare Fig. 5C, D, arrowheads, to Fig. 5G, H, arrowheads). Taken together with our knockdown data, these findings reveal the importance of controlling Cadherin-7 levels in neural crest cells, and ultimately proper neural crest cell-placodal neuron interactions, during trigeminal gangliogenesis.

## DISCUSSION

### Migratory cranial neural crest cells contributing to the trigeminal ganglia express Cadherin-7 throughout early gangliogenesis

The assembly of the cranial trigeminal ganglia requires the coalescence of distinct migratory cell populations, neural crest cells and placodal neurons, that are derived from different tissues (Breau and Schneider-Maunoury, 2015; D’Amico-Martel and Noden, 1983; Hamburger, 1961; Saint-Jeannet and Moody, 2014; Steventon et al., 2014). Cranial neural crest cells leave the neural ectoderm through an epithelial-to-mesenchymal transition and become highly invasive, migrating to the ganglionic anlage (Bronner and Simoes-Costa, 2016; Duband et al., 2015; Gouignard et al., 2018; Simoes-Costa and Bronner, 2015; Taneyhill and Schiffmacher, 2017). During their migration, cranial neural crest cells interact with placodal neurons, which have delaminated from the surface ectoderm and differentiated (Breau and Schneider-Maunoury, 2015; Theveneau et al., 2013). These interactions must be productive to allow for cell-cell adhesion and the eventual correct formation of the cranial ganglia. A prior study noted that N-cadherin expression in trigeminal placodal neurons is critical for mediating proper ganglia condensation (Shiau and Bronner-Fraser, 2009), but molecules important in cranial neural crest cells had yet to be identified. Here, we investigated the role of Cadherin-7 in neural crest cells during trigeminal ganglia assembly. We first documented the spatio-temporal expression pattern of Cadherin-7 protein during early stages of trigeminal gangliogenesis. We observed Cadherin-7 protein exclusively in migratory cranial neural crest cells as early as HH12 (not shown), with no protein present in trigeminal placodal precursors or their neuronal derivatives (Fig. 1). These results now expand upon our prior publication examining the expression and function of adherens junction components in neural crest cells during trigeminal gangliogenesis (αN-catenin in neural crest cells (Wu et al., 2014)). Taken together with these other publications, our new data establish the importance of cadherin-based adhesion during trigeminal ganglia formation, with distinct cadherins expressed in the neural crest cell (Cadherin-7) and placodal neuron (N-cadherin) populations.

### Trigeminal ganglia formation relies upon proper levels of Cadherin-7 in migratory cranial neural crest cells

To explore a function for Cadherin-7 in the cranial neural crest cell population, we undertook molecular perturbation assays to reduce (MO) or elevate (overexpression) Cadherin-7 levels in migratory neural crest cells. MO-mediated knockdown achieved a 50% reduction in Cadherin-7 protein, as assessed by immunoblotting, which, in turn, led to drastic changes in trigeminal ganglia assembly (Figs. 2-3). Migratory neural crest cells depleted for Cadherin-7 still migrated to the ganglionic anlage, but their distribution was altered compared to neural crest cells in control MO-treated embryos. Moreover, trigeminal neurons were also affected at the level of both their morphology and distribution. In many instances these neurons remained round and did not elaborate protrusions characteristic of mature neurons, although they did express Tubb3, indicative of their molecular maturation. This alteration to trigeminal neuron morphology mirrors that observed upon loss of Annexin A6 (Shah et al., 2017) or N-cadherin (Shiau and Bronner-Fraser, 2009) in placodal neurons. Effects on trigeminal neurons upon changes to molecules in neural crest cells are not without precedent, as noted previously upon depletion of αN-catenin in neural crest cells, which leads to an altered trigeminal neuron distribution (noted in transverse sections) and bundling (observed in whole-mount), shown by immunohistochemistry for Islet-1 and Tubb3, respectively (Wu et al., 2014). Moreover, changes in N-cadherin levels in placodal neurons also affect neural crest cells contributing to the cranial ganglia (Shiau and Bronner-Fraser, 2009). Such cell non-autonomous effects are not surprising given that both N-cadherin and Cadherin-7 are transmembrane cell adhesion molecules.

Intriguingly, the trigeminal neurons of embryos possessing lower levels of Cadherin-7 in neural crest cells appear to associate with neural crest cells, but their localization within the anlage was noticeably different than what was observed for control MO-treated embryos. As such, the overall ganglion morphology appeared abnormal. This was particularly apparent in lateral whole-mount views of embryo heads following Cadherin-7 MO electroporation. In these experiments, Cadherin-7-depleted embryos possessed changes in both lobes of the forming trigeminal ganglion, with trigeminal neurons appearing more dispersed and/or less bundled. Collectively, our section and whole embryo results indicate that decreased levels of Cadherin-7 impact overall trigeminal ganglia assembly, likely at the level of both the neural crest cells and placodal neurons.

We next performed the converse experiment in which we overexpressed Cadherin-7 in migratory cranial neural crest cells. With a 200% increase in neural crest Cadherin-7 levels, as evaluated by immunoblotting, we noted, once again, defective trigeminal ganglia assembly, in both section and whole embryo images (Figs. 4-5). The morphology of analyzed trigeminal ganglia was altered upon Cadherin-7 overexpression compared to control pCIG-treated embryos. In these experiments, it is apparent that neural crest cells still migrate to the anlage in the presence of elevated levels of Cadherin-7; however, their distribution is aberrant. Consequently, the location and morphology of trigeminal neurons is also negatively impacted. These neurons, while densely bundled and surrounded by neural crest cells in control embryos, are often times clustered together and exhibit an abnormal shape upon overexpression of Cadherin-7 in neural crest cells. These changes in the distribution of neural crest cells and placodal neurons are also noted in images of the forming trigeminal ganglia in whole embryo heads. Similar to our results in which Cadherin-7 levels are reduced, elevated levels of Cadherin-7 led to a noticeable change in the organization of Tubb3-positive trigeminal neurons. These cells no longer bundle and condense properly with neural crest cells, which instead are observed aggregating with other neural crest cells. Taken together, our knockdown and overexpression results establish a new role for Cadherin-7 in cranial neural crest cells during trigeminal ganglia assembly.

### Neural crest cell-placodal neuron adhesion plays a key role in trigeminal gangliogenesis

Our data indicate that neural crest cells possessing reduced or elevated levels of Cadherin-7 negatively affects trigeminal ganglia assembly, with defects noted in the distribution of neural crest cells and placodal neurons within the ganglionic anlage. Notably, these results cannot be attributed to changes in cell proliferation or cell death within the forming trigeminal ganglia (Supp. Figs. 2, 4). Therefore, our findings further underscore that cell adhesion molecules expressed by neural crest cells and placodal neurons play key roles in regulating ganglia formation. Results published almost a decade ago described the importance of N-cadherin in placodal neurons, including its function in meditating placodal neuron aggregation (Shiau and Bronner-Fraser, 2009). In these experiments, MO-mediated knockdown of N-cadherin in trigeminal placode precursor cells impeded placodal neuron aggregation later in development. Placodal neurons appear more dispersed (evident in section and whole embryo images), much like what we observe for placodal neurons upon Cadherin-7 knockdown in neural crest cells. Moreover, N-cadherin overexpression also resulted in aberrant trigeminal ganglia assembly due to the presence of atypical clusters of placodal neurons, along with an apparent loss of placodal neurons (all noted in whole embryo images). This latter result is intriguing given the comparable placodal neuron phenotypes we observe in whole embryo heads upon Cadherin-7 overexpression in neural crest cells. Unfortunately, effects on the cranial neural crest cell population upon N-cadherin perturbation were not examined in this earlier report. Collectively, our findings reveal that alterations in the levels of cadherins in cranial neural crest cells and trigeminal neurons can severely impact proper ganglia assembly.

Given the observed effect on trigeminal neurons upon changes in neural crest cell Cadherin-7 levels, it is possible neural crest cell corridors do not form entirely correctly in embryos possessing neural crest cells with increased or decreased Cadherin-7 levels. These corridors provide a more permissive substrate (versus the mesoderm) upon which placodal neurons migrate during the formation of the cranial ganglia (Freter et al., 2013). We hypothesize that this could be one mechanism by which changes in neural crest cells affect the distribution and morphology of placodal neurons. Furthermore, elevated levels of Cadherin-7 in neural crest cells could promote increased adhesion between neural crest cells and hinder the ability of these cells to form interactions with placodal neurons. Support for this hypothesis stems from our whole-mount immunohistochemistry images, which show aggregates of neural crest cells after Cadherin-7 overexpression (Fig. 5). Future studies will be necessary to determine whether parameters associated with neural crest cell adhesion and migration (e.g., velocity, directionality) are impacted upon changes in Cadherin-7. In addition, we surmise that alterations in Cadherin-7 levels in neural crest cells could influence N-cadherin distribution and/or levels in placodal neurons, although we have been unable to detect any qualitative changes in N-cadherin by immunohistochemistry upon Cadherin-7 depletion or overexpression. Based on our findings that cadherins are under a high degree of post-translational regulation (e.g., proteolysis: (Schiffmacher et al., 2014)), however, this does not preclude potential changes in placodal neuron adhesion, which will be borne out in future experiments.

In summary, our data provide additional evidence for the importance of properly regulating levels of cadherin proteins during trigeminal ganglia assembly. These findings point to a new role for Cadherin-7 in controlling the formation of the trigeminal ganglia. Altogether, these results further underscore the importance of cadherin-based intercellular interactions that are requisite for cranial gangliogenesis and proper patterning of the vertebrate peripheral nervous system.

## MATERIALS AND METHODS

### Chick embryos

Fertilized chicken eggs (*Gallus gallus*) were obtained from Centurion Poultry (GA) and Moyer’s Chicks, Inc. (PA), and incubated at 37°C in humidified incubators (EggCartons.com, Manchaug, MA, USA). Embryos were staged by the Hamburger-Hamilton (HH) staging method (Hamburger and Hamilton, 1992) or by counting the number of somite pairs (somite stage, ss).

### Cadherin-7 morpholinos and expression constructs

A 3’ lissamine-labeled antisense translation-blocking Cadherin-7 morpholino (MO, 5-ACTCCACTTTGCCCAACTT<u>CAT</u>CTT-3’), or a 5-base pair mismatch Cadherin-7 control MO (5’-AaTCCAaTTTGCCaAAaTT<u>CAT</u>aTT-3’) (start codon underlined, mismatches shown in lowercase), was designed to target the *Cadherin-7* transcript according to the manufacturer’s criteria (GeneTools, LLC). Both MOs were used at a concentration of 500 μM, as described previously (Wu et al., 2014). A DNA construct designed for Cadherin-7 overexpression (pCIG-Cad7), which contains an IRES-GFP cassette to label electroporated cells, was a kind gift from Dr. Marianne Bronner (California Institute of Technology). The control pCIG vector (just the IRES-GFP cassette), or pCIG-Cad7, was used at a concentration of 2.5 μg/μl as in (Wu et al., 2014).

### *In ovo* unilateral electroporations

Unilateral electroporation of the early chick neural tube was conducted to target migratory neural crest cells in the trigeminal ganglionic anlage, as carried out previously (Wu et al., 2014). Briefly, MOs or expression constructs were introduced into premigratory midbrain neural crest cells in developing 3 to 4 somite stage (3-4ss) chick embryos using fine glass needles and filling of the chick neural tube. Platinum electrodes were placed on either side of the embryo, and two 25 V, 25 ms electric pulses were applied across the embryo. Eggs were re-sealed with tape and parafilm, re-incubated for 12 hours, and then imaged *in ovo* around HH12 (prior to embryo turning) using a Zeiss Discovery.V8 stereomicroscope in order to evaluate presence of MOs or expression constructs. After imaging, eggs containing MO- or expression construct-positive embryos were re-sealed and re-incubated for the desired time period prior to harvesting for further experimentation.

### Immunoblotting

Chick embryo neural crest cells were electroporated as described above with either MO or expression constructs. Approximately 35 hours post-electroporation, trigeminal ganglia were excised, pooled, pelleted, flash-frozen in liquid nitrogen, and stored at −80°C until required for immunoblot analysis. Protein lysis, extraction, fraction, and immunoblotting were performed as described previously (Shah et al., 2017; Schiffmacher et al., 2018). Briefly, pellets were thawed on ice and lysed in lysis buffer (50 mM Tris pH 8.0, 150 mM NaCl, 1% IGEPAL CA-630) supplemented with cOmplete protease inhibitor cocktail (Roche, Basel, Switzerland) and 1 mM PMSF for 30 minutes at 4°C with periodic mixing. Soluble fractions were collected following centrifugation at max g for 15 minutes at 4°C, and protein concentration was quantified by Bradford assay (Thermo Fisher Scientific, Rockford, IL, USA). Equivalent amounts of protein per sample were processed by SDS-PAGE (10% Mini-Protean TGX gel, BioRad #456-1034) and then transferred to 0.45μm BioTrace PVDF membrane (Pall, Port Washington, NY) via the iBlot transfer stack system (iBlot 2 Dry Blotting system, Life Technology # IB21001) according to the manufacturer’s guidelines. Primary antibodies used for immunoblotting were Cadherin-7 (Developmental Studies Hybridoma Bank (DSHB), clone CCD7-1, 1:150) and β-actin (Santa Cruz Biotechnology sc-47778, 1:1000). Immunoblot images for figures were gamma-modified and processed using Adobe Photoshop CC 2015.5 (Adobe Systems, San Jose, CA, USA). Immunoblot band volumes (intensities) were calculated from unmodified immunoblot images using Image Lab software (Bio-Rad, Hercules, CA, USA), and relative protein levels were determined by normalizing the volumes of Cadherin-7 bands to those of β-actin. Differences in the amount of Cadherin-7 were assessed by comparing normalized ratios between either control MO- and Cadherin-7 MO-treated samples, or pCIG- and pCIG-Cad7-treated samples, with the control MO- and pCIG-treated samples set to one.

### Immunohistochemistry and TUNEL assay

Embryos were collected at the designated stages for wild-type or post-electroporation immunohistochemistry. Detection of various proteins was performed in whole-mount following overnight fixation in 4% PFA, or on 14μm transverse sections following 4% PFA fixation, gelatin embedding, and cryostat sectioning as described previously (Shah et al., 2017; Wu et al., 2014). All primary and secondary antibodies were diluted in 1X Phosphate-buffered saline + 0.1% Triton X-100 (PBSTX) + 5% sheep serum. The following antibodies and dilutions were used for immunohistochemistry: Cadherin-7 (DSHB, clone CCD7-1, 1:100); N-cadherin (DSHB, clone MNCD2, 1:200); HNK-1 (DSHB, clone 3H5, 1:100;); Tubb3 (Abcam 2G10, ab78078, 1:500); Annexin A6 (Abnova, PAB18085, 1:100); GFP (Abcam, ab6662, 1:300); and phospho-histone H3 (Millipore, 1:200). The following secondary antibodies were used at 1:200-1:500 dilutions: goat anti-mouse IgG (Life Technologies, Cadherin-7); goat anti-rat IgG (Life Technologies, N-cadherin); goat anti-mouse IgM (Life Technologies, HNK-1); goat anti-mouse IgG_2a_ (Southern Biotech, Tubb3); and goat anti-rabbit IgG (Life Technologies, Annexin A6 and phospho-histone H3). Sections were stained with 4′,6-diamidino-2-phenylindole (DAPI) to mark cell nuclei using DAPI-containing mounting media (Fluoromount G, Southern Biotech). A TUNEL assay (Roche, TMR red and fluorescein) was performed on 4% PFA-fixed, cryopreserved sections to detect apoptotic cells as described previously (Shah et al., 2017; Wu et al., 2014) followed by mounting of slides with DAPI-containing media as outlined above.

### Confocal Imaging

For all experiments, images of at least five serial transverse sections through a minimum of eight embryos (unless indicated otherwise), or of a minimum of seven embryo heads (unless indicated otherwise), were acquired with the LSM Zeiss 800 confocal microscope with Airyscan detection (Carl Zeiss Microscopy, Thornwood, NY, USA) at 20X or 5X magnification, respectively. To acquire images of the trigeminal ganglion in the chick head, embryos were mounted on viewing slides, and a lateral view of the chick head containing the forming trigeminal ganglion was captured. Where possible, the laser power, gain, and offset were kept consistent for the different channels throughout all experiments. Image processing was conducted with the Zen Blue software (Carl Zeiss Microscopy) and Adobe Photoshop CC 2015.5.

### Quantification and statistical analysis

To analyze the effect of Cadherin-7 knockdown or overexpression on cell proliferation and cell death, phospho-histone H3- and TUNEL-positive cells were counted following immunohistochemistry (or TUNEL assay) using the Adobe Photoshop count tool. Cells were counted within the region of the forming trigeminal ganglion in a minimum of five serial transverse sections taken from at least three electroporated embryos per treatment, on both the experimentally-treated and contralateral control sides of the section. Cell counts were then compared within embryo treatment groups. All results are reported as the average number of phospho-histone H3- or TUNEL-positive cells, plus or minus the standard error of the mean, and were analyzed with an unpaired Student’s t test to establish statistical significance as carried out previously (Shah et al., 2017; Wu et al., 2014).

## ACKNOWLEDGMENTS

We thank Ms. Vinona Muralidaran, Ms. Reethika Maddineni, and Ms. Julie Ren for excellent technical assistance. We also thank Dr. Marianne Bronner (California Institute of Technology) for the Cadherin-7 expression construct (pCIG-Cad7). The authors declare no competing financial interests. This work was supported by a grant to L.A.T. (NIH R01DE024217).

## SUPPLEMENTAL FIGURE LEGENDS

**Supplemental Figure 1.**
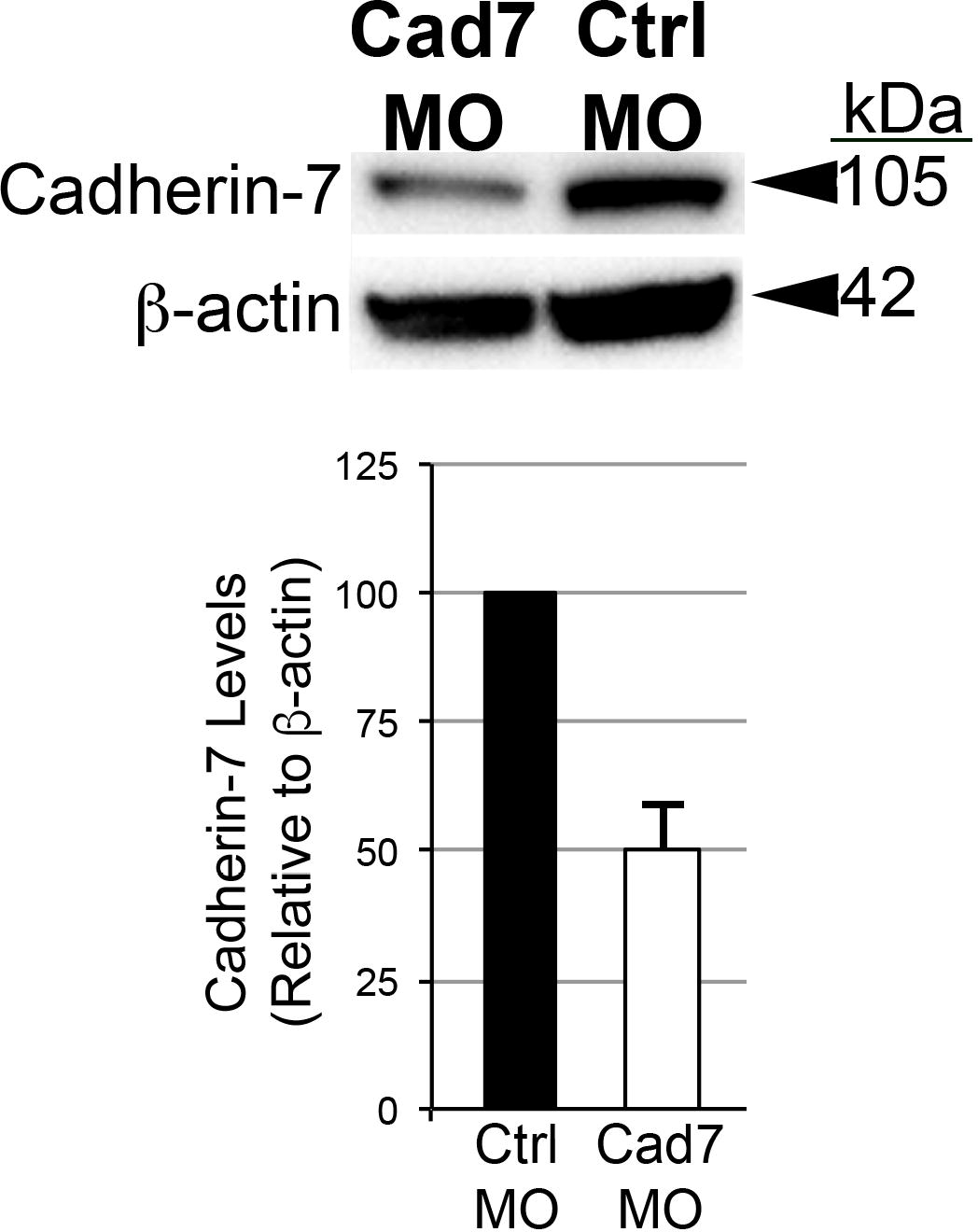
Knockdown of Cadherin-7 with a translation-blocking morpholino antisense oligonucleotide targeting *Cadherin-7* effectively reduces Cadherin-7 protein in migratory neural crest cells contributing to the trigeminal ganglia. Premigratory neural crest cells were electroporated at the 2-3ss with either the Cadherin-7 morpholino (Cad7 MO) to allow for depletion of Cadherin-7 protein in migratory neural crest cells, or a 5 bp mismatch Cadherin-7 control MO (Ctrl MO). Embryos were re-incubated to HH15-17 after which time the trigeminal ganglion-forming region on the electroporated side of the embryo was dissected out of the embryo and pooled for lysate preparation. Immunoblotting for Cadherin-7 and β-actin (control) was performed as in (Shah et al., 2017), with a representative immunoblot shown. Knockdown efficiency was assessed as previously described (Shah et al., 2017), with graph revealing results of immunoblot analysis as determined by normalizing Cadherin-7 to β-actin and calculating the reduction in this normalized ratio from that obtained for the control MO-treated lysate (arbitrarily set to 1, n = 2). The mean and standard error of the mean are shown. A 50% knockdown in Cadherin-7 protein levels is noted in the Cadherin-7 MO-treated lysate compared to the control MO-treated lysate.

**Supplemental Figure 2.**
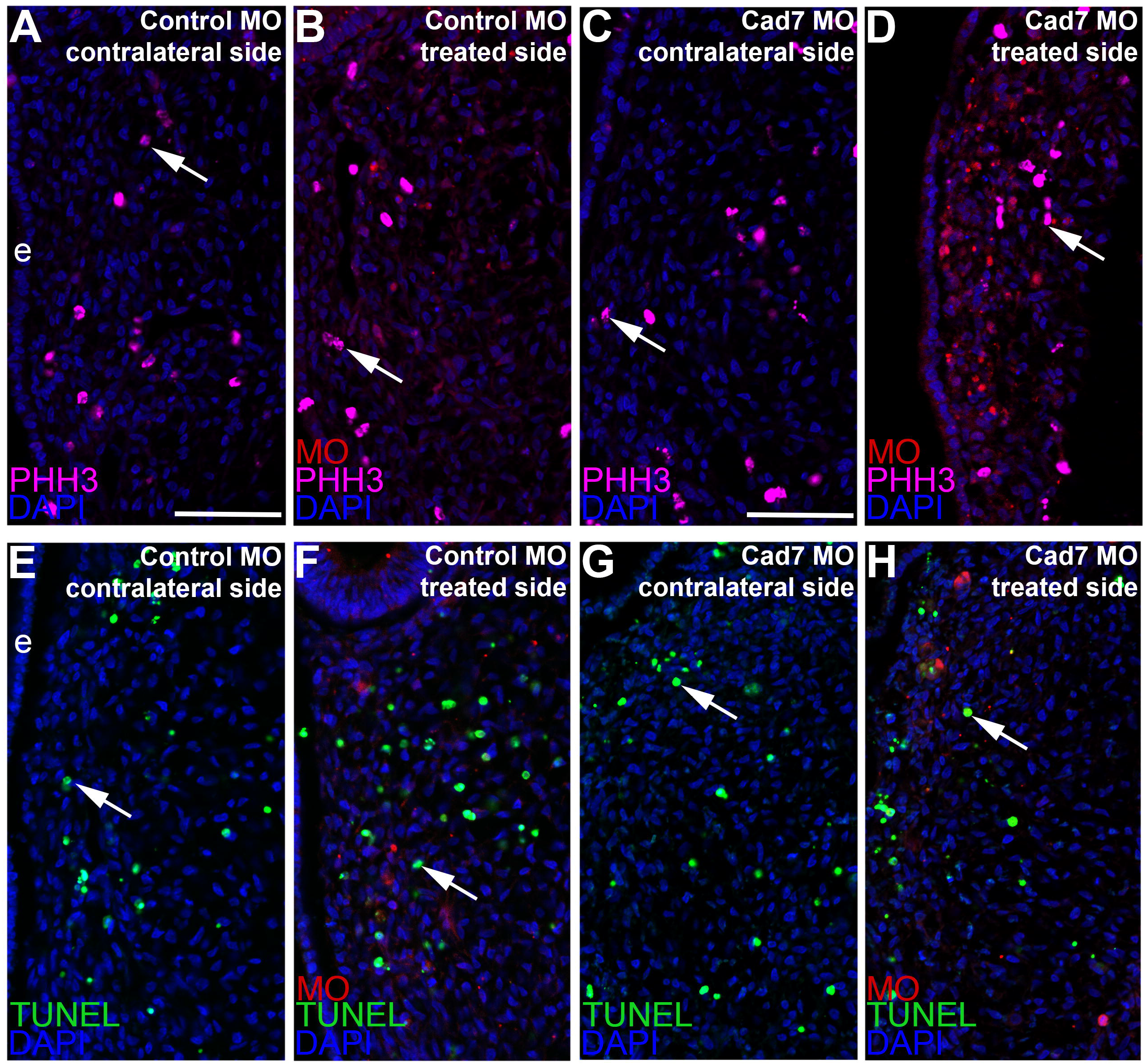
Electroporation of either the control or Cadherin-7 morpholino does not alter cell proliferation or cell death in the trigeminal ganglionic anlage. Representative transverse sections taken at the axial level of the the forming trigeminal ganglia after electroporation of a 5 bp mismatch control Cadherin-7 morpholino (Control MO: A, B, E, F) or Cadherin-7 morpholino (Cad7 MO: C, D, G, H) into premigratory neural crest cells at the 3ss followed by immunohistochemistry for phospho-histone H3 (PHH3, A-D) or TUNEL (E-H). Contralateral (A, C, E, G) and morpholino-treated (B, D, F, H) sides are shown to provide a means of comparison. Arrows indicate PHH3 (A-D)- and TUNEL (E-H)-positive nuclei, with a comparable number noted in the presence of either morpholino relative to the contralateral control side of the electroprated embryo. DAPI (blue) labels cell nuclei. Ectoderm (e) is oriented to the left within each image panel and may not be visible in the field of view for some images. Scale bar in (A) is 67μm and applies to (B, E, F), while scale bar in (C) is 50μm and applies to (D, G, H).

**Supplemental Figure 3.**
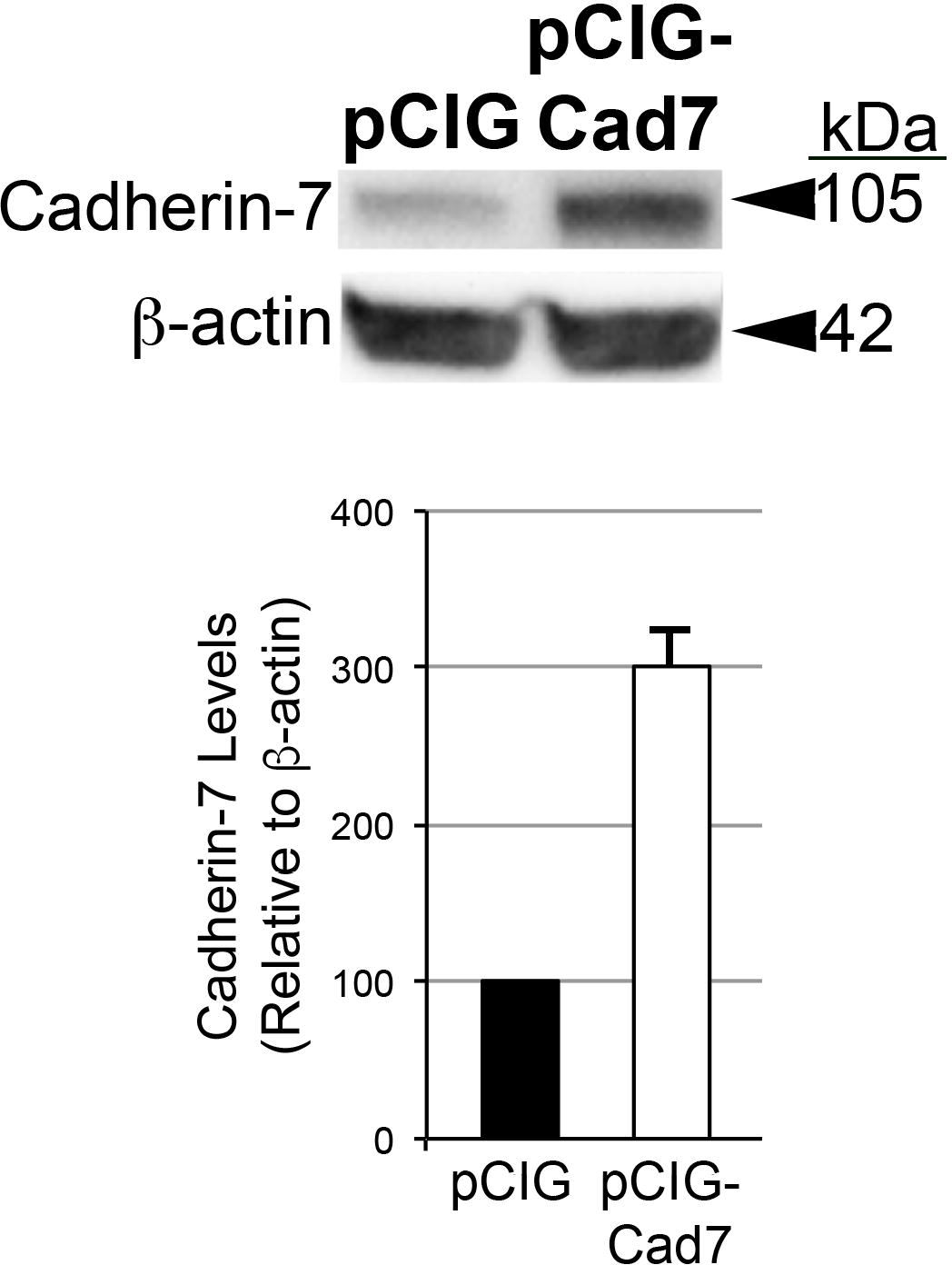
Overexpression of Cadherin-7 effectively increases Cadherin-7 protein in neural crest cells contributing to the trigeminal ganglia. Premigratory neural crest cells were electroporated at the 2-3ss with either a Cadherin-7 expression construct (pCIG-Cad7) to allow for overexpression of Cadherin-7 protein in migratory neural crest cells, or the control vector (pCIG). Embryos were re-incubated to HH15-17 after which time the trigeminal ganglion-forming region on the electroporated side of the embryo was dissected out of the embryo and pooled for lysate preparation. Immunoblotting for Cadherin-7 and β-actin (control) was performed as in (Shah et al., 2017), with a representative immunoblot shown. Overexpression efficiency was assessed as previously described (Shah et al., 2017), with graph indicating results of immunoblot analysis as determined by normalizing Cadherin-7 to β-actin and calculating the increase in this normalized ratio from that obtained for the pCIG-treated lysate (arbitrarily set to 1, n = 2). The mean and standard error of the mean are shown. A 200% increase in Cadherin-7 protein levels is noted in the pCIG-Cad7-treated lysate compared to the control pCIG-treated lysate.

**Supplemental Figure 4.**
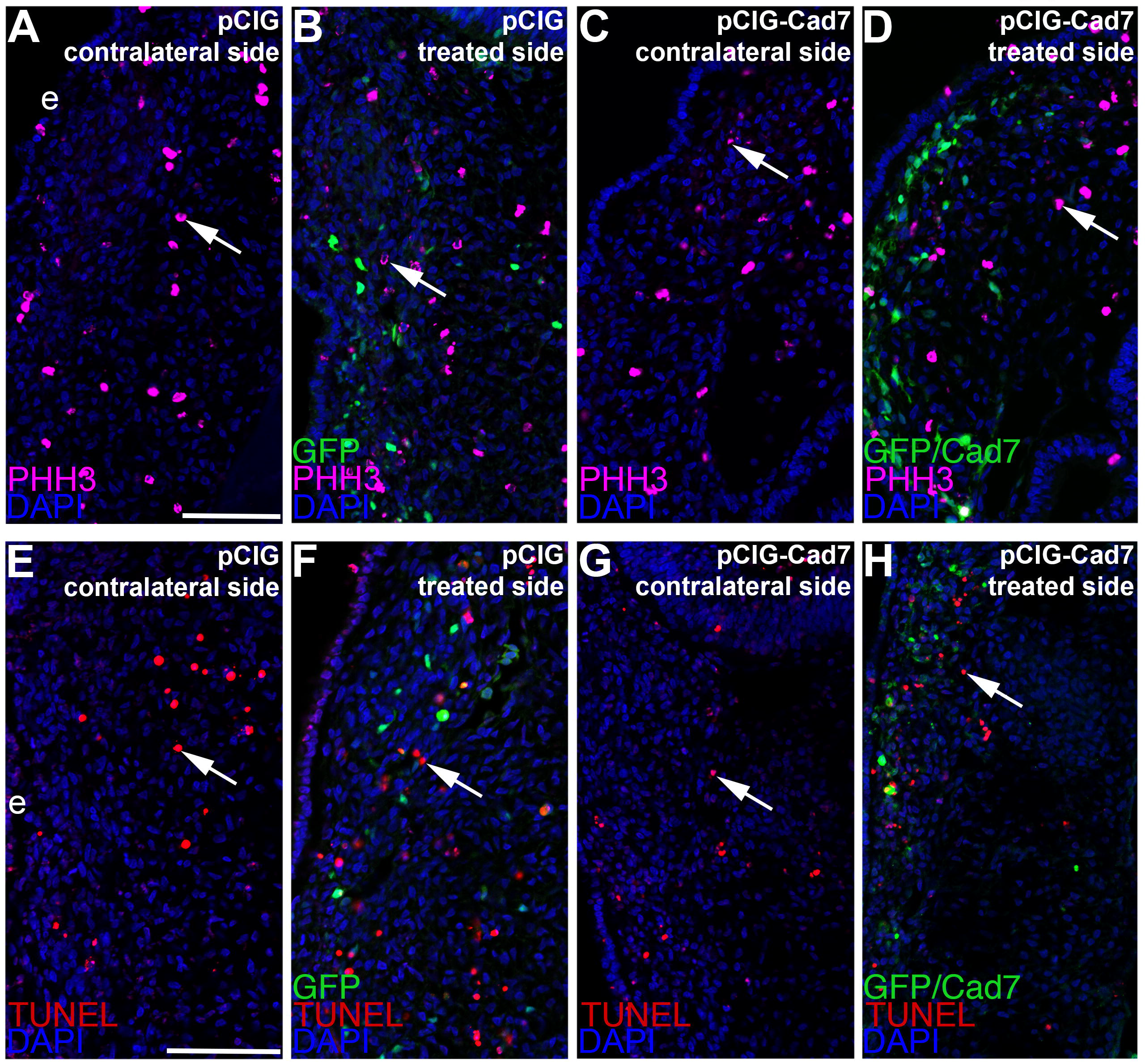
Electroporation of expression constructs does not alter cell death or cell proliferation in the trigeminal ganglionic anlage. Representative transverse sections taken at the axial level of the forming trigeminal ganglia after electroporation of the pCIG control vector (pCIG: A, B, E, F) or pCIG-Cadherin-7 vector (pCIG-Cad7: C, D, G, H) into premigratory neural crest cells at the 3ss followed by immunohistochemistry for phospho-histone H3 (PHH3, A-D) or TUNEL (E-H). Contralateral (A, C, E, G) and expression vector-treated (B, D, F, H) sides are shown to provide a means of comparison. Arrows indicate PHH3 (A-D)- and TUNEL (E-H)-positive nuclei, with a comparable number noted in the presence of either expression construct relative to the contralateral control side of the electroprated embryo. DAPI (blue) labels cell nuclei. Ectoderm (e) is oriented to the left within each image panel. Scale bar in (A) is 60μm and applies to (B-D), while scale bar in (E) is 60μm and is applies to (F-H).

